# Simple Electroporation of *Chlamydomonas reinhardtii* Strains with an Intact Cell Wall

**DOI:** 10.64898/2026.04.30.721989

**Authors:** Maximilian Meßmer, Félix de Carpentier, Ezekiel Lam, Meggie Hong, Setsuko Wakao, Michael Schroda, Krishna K. Niyogi

## Abstract

*Chlamydomonas reinhardtii* is a model green alga extensively used to study photosynthesis and cilia using molecular biology and genetics. Electroporation is a very common technique to transform DNA into the nuclear genome, which is essential to generate mutant collections and express transgenes. Here, we describe a simple, fast, and efficient protocol to transform strains with an intact cell wall. It achieves a good transformation efficiency without cell wall digestion or use of commercial kits and is compatible with the widely available Gene Pulser electroporation system.

**Key features:**

- High transformation efficiency of *Chlamydomonas reinhardtii* strains with an intact cell wall.
- Faster than currently available electroporation protocols.

## Background

*Chlamydomonas reinhardtii* is a model green alga amendable to genetic engineering. Electroporation is a common technique to randomly integrate DNA into the nuclear genome through non-homologous end joining repair [7]. Several protocols have been developed, but they often require cell-wall-less strains, digestion of the cell wall, specialized electroporators (*e*.*g*. NEPA21, Nepagene) [13], or a commercial kit (*i*.*e*. MAX Efficiency Transformation Reagent for Algae kit, Invitrogen). Here, we describe a simple, fast, and efficient method to transform *Chlamydomonas reinhardtii* strains with an intact cell wall. Compared to previously published electroporation protocols by Crozet et al. [2,3] and Onishi and Pringle [12], we simplified the procedure by removing the incubation in the cold (20 to 30 min), allowing the electroporation part of our protocol (section C) to take around 30 minutes. We used a CHES pH 9.25 electroporation buffer [12], which has been reported to yield a high number of transformants and to be efficient with several transgenes and strains including the ones with an intact cell wall [1,4,10,12]. Our protocol is suitable for generation of randomly mutagenized mutant collections and transgenic approaches (*e*.*g*. complementation of mutants or protein localization) but not for delivery of Cas9 ribonucleoprotein complexes.

## Materials and reagents

### Biological materials

1. *Chlamydomonas reinhardtii*, tested strains: CC-4051 (4A^-^), CC-5101 (T222^+^), CC-4425 (D66), CC-124, CC-4533, CC-5325 (available at the Chlamydomonas Resource Center, www.chlamycollection.org). Does not work with true cell-wall-less strains like UVM4, for which glass bead transformation is recommended [11].
2. Expression plasmid for *Chlamydomonas reinhardtii*. This protocol does not cover plasmid design and cloning, but we recommend using MoClo vectors [2,11] (available at the Chlamydomonas Resource Center, www.chlamycollection.org/product/moclo-toolkit).

### Reagents

1. Tris base (tris(hydroxymethyl)aminomethane, Fisher Scientific, BP152-1)
2. Potassium phosphate dibasic (K_2_HPO_4_, Fisher Scientific, BP363-500)
3. Potassium phosphate monobasic (KH_2_PO_4_, VWR, EMD-PX1565-1)
4. Ammonium chloride (NH_4_Cl, VWR, BDH9208-500G)
5. Magnesium sulphate heptahydrate (MgSO_4_.7H_2_O, Fisher Scientific, M63-500)
6. Calcium chloride dihydrate (CaCl_2_.2H_2_O, Fisher Scientific, C79-500)
7. EDTA-Na_2_ (Ethylenediaminetetraacetic acid disodium salt dihydrate, Sigma-Aldrich, E4884)
8. Potassium hydroxide (KOH, Sigma-Aldrich, P6310)
9. Ammonium molybdate tetrahydrate ((NH_4_)_6_Mo_7_O_24_, Fisher Scientific, A674-500)
10. Zinc sulfate heptahydrate (ZnSO_4_.7H_2_O, Sigma-Aldrich, Z4750)
11. Manganese(II) chloride tetrahydrate (MnCl_2_.4H_2_O, Sigma-Aldrich, M3634)
12. Iron(III) chloride hexahydrate (FeCl_3_.6H_2_O, Sigma-Aldrich, F2877)
13. Sodium carbonate (Na_2_CO_3_, Sigma-Aldrich, BP357-1)
14. Copper(II) chloride dihydrate (CuCl_2_.2H_2_O, Sigma-Aldrich, 307483)
15. Glacial acetic acid (CH_3_COOH, VWR, EMD-AX0073-9)
16. Hydrochloric acid (HCl, Fisher Scientific, A144SI-212) 17.
17. Agar (VWR, 90000-762)
18. Blasticidin S hydrochloride (Fisher Scientific, 50-712-729)
19. Zeocin (Thermo Fisher Scientific, R25001)
20. Hygromycin B (Thermo Fisher Scientific, 10687010)
21. Kanamycin monosulfate (TCI, K0047)
22. Paromomycin sulfate (Sigma-Aldrich, P5057)
23. Spectinomycin dihydrochloride pentahydrate (Sigma-Aldrich, S4014)
24. Nourseothricin sulfate (GoldBio, N-500)
25. CHES (*N*-cyclohexyl-2-aminoethanesulfonic acid, Sigma-Aldrich, C2885)
26. Sucrose (Millipore, 573113)
27. Sorbitol (Thermo Fisher Scientific, 036404.36)

### Recipes

#### 1. 1M Tris base

1. Weigh 121.14 g Tris.
2. Dissolve in 800 ml of distilled water.
3. Adjust the volume to 1000 ml with distilled water.

#### 2. Phosphate Buffer II

1. Weigh:
  - 10.8 g K_2_HPO_4_
  - 5.6 g KH_2_PO_4_
2. Dissolve one by one in 80 ml of distilled water.
3. Adjust the volume to 100 ml with distilled water.

#### 3. Solution A

1. Weigh:
  - 20.0 g NH_4_Cl
  - 5.0 g MgSO_4_.7H_2_O
  - 2.5 g CaCl_2_.2H_2_O
2. Dissolve one by one in 400 ml of distilled water.
3. Adjust the volume to 500 ml with distilled water.

#### 4. Kropat’s stock solutions

Kropat’s trace elements solutions preparation is based on the following protocol www.chlamycollection.org/content/uploads/2022/05/Kropats-Trace-Elements-safety-update.pdf [9]. Selenite is omitted as it is not necessary for this protocol. If sodium selenite is used, it should be handled with care as it is toxic.

1. 125 mM EDTA-Na_2_ stock solution
  - Dissolve 11.63 g of EDTA-Na_2_ in 250 ml of distilled water.
  - Adjust to pH8.0 with KOH pellets.
  - Adjust the volume to 300 ml with distilled water.
2. 285 μM (NH_4_)_6_Mo_7_O_24_
  - Dissolve 88.0 mg of (NH_4_)_6_Mo_7_O_24_ is 200 mL of distilled water.
  - Adjust the volume to 250 ml with distilled water.

#### 5. Kropat’s trace elements solutions (1000x-concentrated)

1. 25 mM EDTA-Na_2_
  - Mix 50 ml of 125 mM EDTA-Na_2_ and 200 ml of distilled water.
2. 28.5 μM (NH_4_)_6_Mo_7_O_24_
  - Mix 25 ml of 285 μM (NH_4_)_6_Mo_7_O_24_ and 225 ml of distilled water.
3. Zn-EDTA (2.5 mM ZnSO_4_, 2.75 mM EDTA-Na_2_)
  - Dissolve 180.0 mg of ZnSO_4_.7H_2_O in 200 ml of distilled water.
  - Add 5.5 ml of 125 mM EDTA-Na_2_.
  - Adjust the volume to 250 ml with distilled water.
4. Mn-EDTA (6 mM MnCl_2_, 6 mM EDTA-Na_2_)
  - Dissolve 297.0 mg of MnCl_2_.4H_2_O in 200 ml of distilled water.
  - Add 12 ml of 125 mM EDTA-Na_2_.
  - Adjust the volume to 250 ml with distilled water.
5. Fe-EDTA (20 mL FeCl_3_, 22 mM EDTA-Na_2_, 22 mM Na_2_CO_3_)
  - Dissolve 2.05 g of EDTA-Na_2_ and 0.58 g of Na_2_CO_3_ in 200 ml of distilled water.
  - Add and dissolve 1.35 g of FeCl_3_.6H_2_O.
  - Adjust the volume to 250 ml with distilled water.
6. Cu-EDTA (2 mM CuCl_2_, 2 mM EDTA-Na_2_)
  - Dissolve 85.0 mg of CuCl_2_.2H_2_O in 200 ml in distilled water.
  - Add 4 ml 125 mM EDTA-Na_2_.
  - Adjust the volume to 250 ml with distilled water.

#### 6. Liquid TAP

1. Add the following solutions to 900 ml of distilled water.
  - 20 ml 1M Tris base
  - 1 ml Phosphate Buffer II
  - 10 ml Solution A
  - 1 ml of **each** Kropat’s trace elements solution: 25 mM EDTA-Na_2_, 28.5 μM (NH_4_)_6_Mo_7_O_24_, Zn-EDTA, Mn-EDTA, Fe-EDTA, and Cu-EDTA.
  - 1 ml of glacial acetic acid
2. Adjust pH to 7.0 with concentrated HCl.
3. Adjust the volume to 1000 ml with distilled water.
4. Autoclave for 30 to 40 min.

#### 7. TAP plates with antibiotics

1. Prepare TAP as in recipe 5 (without the autoclaving step).
2. Add 15 g of agar per liter.
3. Autoclave for 30 to 40 min.
4. Mix well and let the solution reach a temperature of 40 to 50°C.
5. Add relevant antibiotic at the following final concentration: blasticidin S (50 mg/l), zeocin (15 mg/l), hygromycin B (20 mg/l), kanamycin (50 mg/l), paromomycin (20 mg/l), spectinomycin (100 mg/l), or nourseothricin (10 mg/l) [3,14].
6. Mix well and pour plates.
7. Store at 4-8°C for up to 2 months.

#### 8. CHES buffer (10 mM CHES pH 9.25, 40 mM sucrose, 10 mM sorbitol)

1. Weigh:
  - 1 g CHES
  - 6.84 g sucrose
  - 0.91 g sorbitol
2. Dissolve one by one in 400 ml of distilled water.
3. Adjust pH to 9.25.
4. Bring to 500 ml with distilled water.
5. Filter sterilize.

#### 9. TS40 (40 mM sucrose in TAP)

1. Dissolve 6.84 g sucrose in 500 ml TAP.
2. Filter sterilize.

### Laboratory supplies

1. Gel extraction kit (*e*.*g*. Monarch Spin DNA, New England Biolabs, T1120S)
2. Bottle top 0.45 μm filers (*e*.*g*. Nalgene Rapid flow, Thermo Scientific, 565-0010)
3. 4 mm-gapped electroporation cuvettes (USA Scientific, 9104-6050)
4. Parafilm or breathable tape (3M, Vent Tape 394)

## Equipment

1. Gene Pulser with Pulse Controller (Bio-Rad)
2. NanoDrop spectrophotometer (ThermoFisher Scientific)
3. Hemocytometer or cell counter (*e*.*g*. Multisizer 4e Coulter Counter, Beckman Coulter)
4. Growth chamber with light and temperature control (*e*.*g*. Percival, Conviron, BioChamber) and a rotary shaker. A simpler culture setup such as hanging fluorescent bulbs over a shaker at room temperature has yielded similar results.
5. Standard laboratory equipment (*e*.*g*. pipettes, centrifuge, microcentrifuge, laminar flow hood, glass beakers, 15- and 50-ml sterile tubes, *etc*.)

## Procedure

### A. Purification of the DNA cassette

1. Digest a sufficient quantity of plasmid DNA (typically 5 to 10 µg) with appropriate restriction enzymes (typically *Bbs*I or *Bsa*I for MoClo plasmids [2,11]) to isolate the desired cassette. If the size of the cassette is close to that of the vector backbone fragment, additionally cut the latter with another restriction enzyme to ensure good separation. It is also possible to linearize the plasmid without purifying the expression cassette on gel [1].
2. Incubate the reaction for 3 h to overnight at 37°C.
3. Separate the cassette by agarose gel electrophoresis and isolate it by gel purification.
4. Measure the DNA concentration with a NanoDrop spectrophotometer. Typical recovery yield is 60-80%.

### B. Chlamydomonas culture and harvesting

1. From this point on, all steps must be performed under sterile conditions, for example in a laminar flow hood.
2. Inoculate *Chlamydomonas reinhardtii* culture in liquid TAP. Adjust the volume depending on the number of transformations, the minimum is 25 mL per transformation if cells are harvested at 2 x 10^6^ cells/ml.
3. Incubate the culture at 25°C with shaking (∼120 rpm) and constant light (50-100 μmol photons m^-2^ s^-1^) until it reaches a concentration of 2-5 x 10^6^ cells/ml which takes 2 to 3 days. Cell concentration can be measured with a hemocytometer or a cell counter. Using optical density has not specifically been tested but may be used following the methods developed in [6]. It is recommended to maintain the culture under 5 x 10^6^ cells/ml by dilution when necessary. This protocol works with stationary phase cultures as well, but efficiency might be slightly reduced.

### C. Electroporation and recovery

1. In 50 ml tubes, harvest 50 x 10^6^ cells per transformation by centrifugation (2500 x *g*, room temperature, 5 min) - calculate the volume to harvest as follows: V = 5 x 10^7^ / concentration. For example, if the culture concentration is 2 x 10^6^ cells/ml, the volume is V = 50 x 10^6^ / 2 x 10^6^ = 25 mL per transformation.
2. Remove all the supernatant and resuspend the cell pellet in CHES buffer to a final concentration of 2 x 10^8^ cells/ml.
3. In a 4 mm-gapped electroporation cuvette, mix 250 μl of the cell suspension with 50-500 ng of DNA cassette. 200 ng of DNA is usually sufficient, but the quantity may need to be adjusted depending on the expression cassette. Several samples can be prepared at the same time as incubation time has little effect on the transformation efficiency.
4. Right before electroporation, gently flick the cuvette to resuspend the cells. If bubbles are present, tap the cuvette on the bench to eliminate them.
5. Proceed to the electroporation at 600 V, 25 µF, and 1000 Ω. Electroporation also works with Bio-Rad systems that lack resistance control. *Note: The measured pulse time has been between 10 and 15 ms for successful transformations*.
6. Transfer the cells to a 15 mL tube filled with approximately 10 ml TS40 and rinse the cuvette twice with 500 μl TS40 to recover all the cells.
7. Incubate the tubes at 25°C and a light intensity of ∼50 μmol photons m^-2^ s^-1^ horizontally on a rotary shaker at ∼120 rpm (or in a tube rotator at moderate speed) for 12 to 16 h.

*Note: Do not skip this recovery step, as it is essential for the cells to express the resistance gene*.

### D. Selection of transformants

1. Harvest cells by centrifugation (2500 x *g*, room temperature, 5 min).
2. Pour off almost all the supernatant, leaving a volume of ∼300 to 500 μl.
3. Resuspend the cell pellet in the remaining liquid, spread on TAP agar plates supplemented with the relevant antibiotic, and dry.
4. Seal the plates with parafilm or breathable tape and incubate at 25°C and 50-100 μmol photons m^-2^ s^-1^.
5. Colonies should appear after 4 to 6 days (Figure 1).

**Figure 1.**
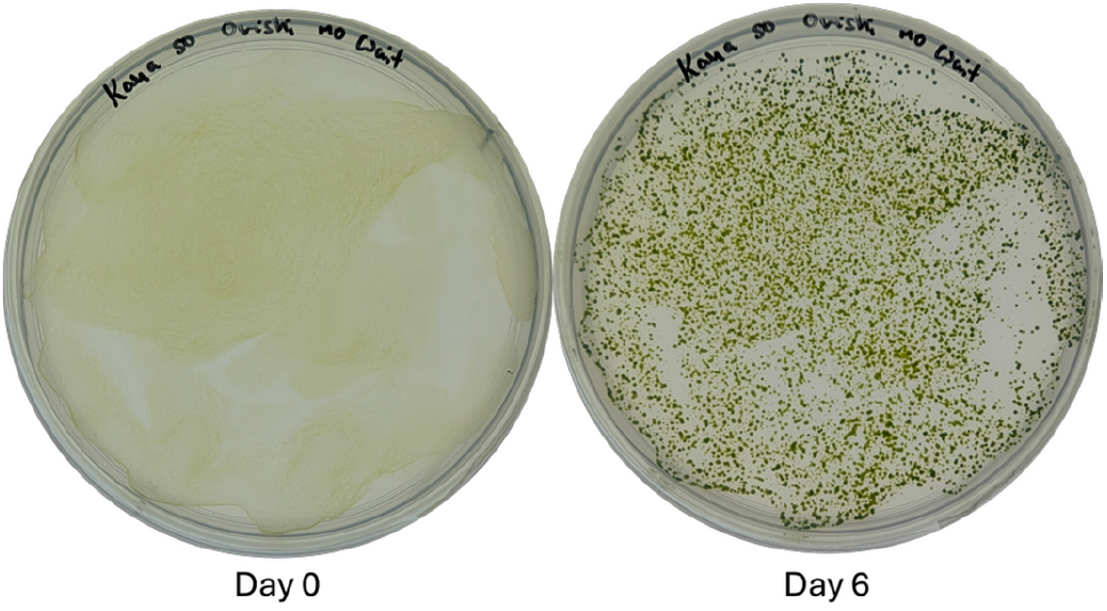
A typical successful electroporation. Pictures were taken 0 and 6 days after plating. See Figure 2 legend for more details.

## Validation of protocol

This protocol or a very similar one was successfully used in the following publications [1,4,10,12]. Here, it was further validated by comparing it with the ‘Crozet’ protocol [3] and the MAX Efficiency commercial kit. Our protocol produced comparable results in terms of number of transformants or transgene expression, as estimated here by mVenus fluorescence (Figure 2). Transformants characterization depends on the goal of the experiment, it may include detection or localizing the insertion locus by PCR [3,8], or detection of the produced protein [1].

**Figure 2.**
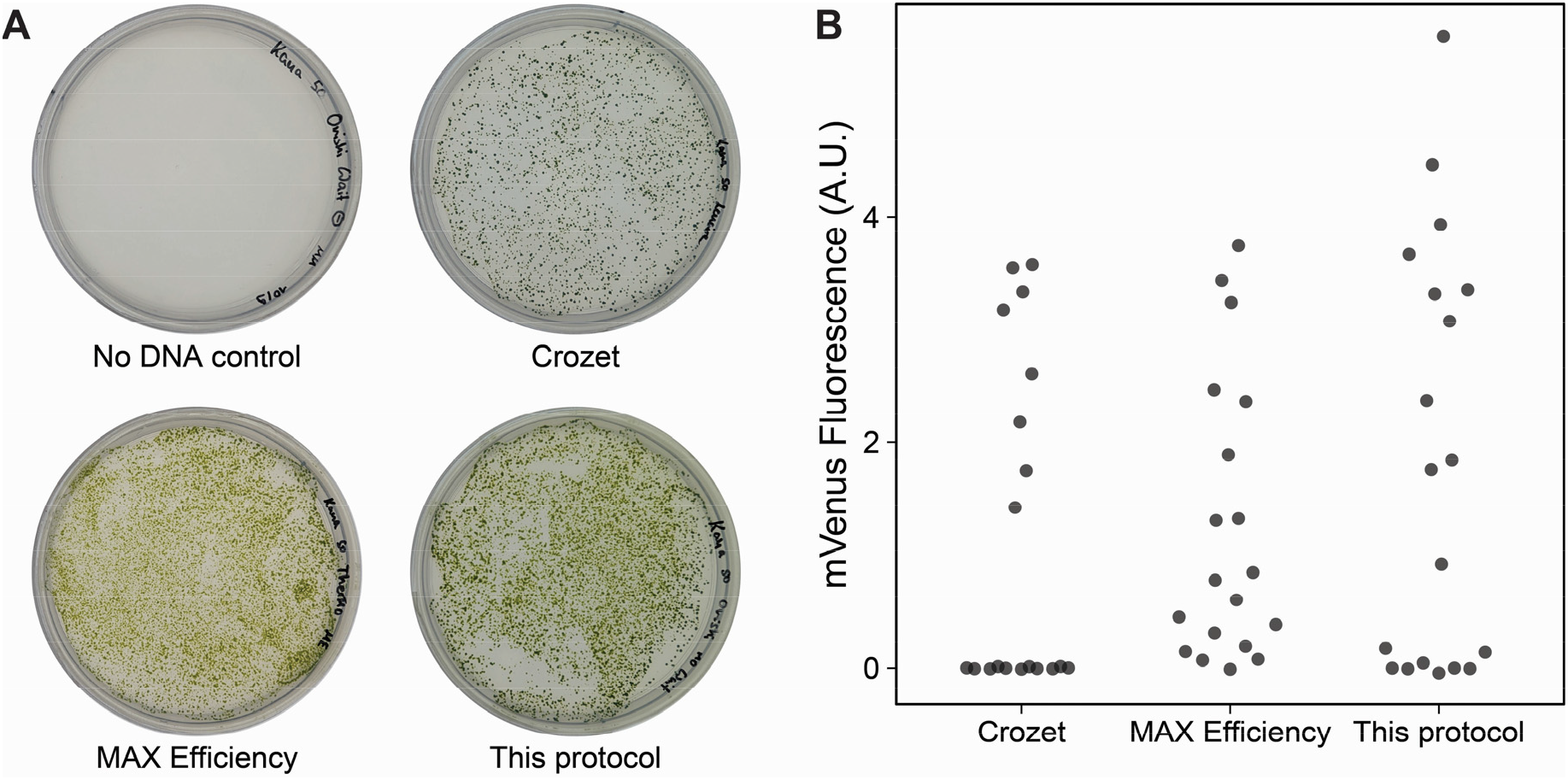
Validation of the electroporation protocol in a wild type strain with an intact cell wall. (A) The plasmid pFCM-010 (pAR-*NPTII*-tRbcS2-pAβSAP-*mVenus*-tPsaD, Dataset S1) was built with the Chlamydomonas MoClo toolkit [2,5]. 500 ng of purified cassette were transformed into the 4A^-^ wild type (with an intact cell wall) with our simplified protocol, the ‘Crozet’ protocol [2,3], or the MAX Efficiency Transformation Reagent for Algae kit (Invitrogen) and selected on TAP+kanamycin plates. The transformation efficiency of our protocol was estimated at 13.3 x 10^3^ transformants/μg of DNA or 134 transformant/10^6^ cells, and the one of the ‘Crozet’ protocol was of 7.0 x 10^3^ transformants/μg of DNA or 70 transformants/10^6^ cells. Note that the number of transformants may depend on the expression cassette. Pictures were taken after 6 days. (B) Fluorescence of mVenus in individual transformants generated with the three protocols. 96 well plates containing 200 μl of cell culture per well were measured with a Tecan Infinite M1000 PRO plate reader (excitation: 505 nm; emission: 530 nm) and normalized with chlorophyll fluorescence (excitation: 430 nm; emission: 655nm).

### Troubleshooting

Problem: Low transformation efficiency.

Possible cause: Saturated or unhealthy culture.

Solution: Repeat the experiment and check that the preculture contains mostly single cells. If the strain is motile, cells should be

swimming. Also check that the culture is not contaminated by plating cells on Luria broth.

## Supplementary information

The following supporting information can be downloaded here (link available when this protocol is published online):

1. Dataset S1. Plasmid map of pFCM-010

## Acknowledgments

Conceptualization, M.M., F.C.; Investigation, M.M., F.C., E.L., M.H.; Data Analysis, M.M., F.C., S.W., M.S., K.K.N.; Writing-Original Draft, M.M., F.C.; Writing-Review & Editing, M.M., F.C., E.L., M.H., S.W., M.S., K.K.N.; Funding acquisition, M.S., K.K.N.; Supervision, F.C., S.W., K.K.N.

We acknowledge Dr. Masayuki Onishi for developing the original version of the protocol and Dr. Olivier D. Caspari for providing it to us. This work was supported by the Howard Hughes Medical Institute, the BioComp 4.0 program (RPTU), and UC Berkeley Sponsored Projects for Undergraduate Research (SPUR) program. K.K.N. is an investigator of the Howard Hughes Medical Institute and S.W. is supported by the U.S. Department of Energy, Office of Science, through the Photosynthetic Systems program in the Office of Basic Energy Sciences.

